# A tumor suppressive long noncoding RNA, DRAIC, inhibits protein translation and induces autophagy by activating AMPK

**DOI:** 10.1101/2021.06.04.447165

**Authors:** Shekhar Saha, Ying Zhang, Briana Wilson, Roger Abounader, Anindya Dutta

**Affiliations:** Department of Biochemistry and Molecular Genetics, University of Virginia School of Medicine, Charlottesville, Virginia, USA 22901; Department of Genetics, University of Alabama, Birmingham, Alabama, USA 35233; Department of Microbiology, Immunology and Cancer Biology, University of Virginia School of Medicine, Charlottesville, Virginia, USA 22901; Cancer Center, University of Virginia, Charlottesville, VA, USA

**Keywords:** DRAIC lncRNA, AMPK, mTORC1, protein-translation, autophagy

## Abstract

LncRNAs are long RNA transcripts that do not code for proteins and that have been shown to play a major role in cellular processes through diverse mechanisms. DRAIC, a lncRNA which is downregulated in castration-resistant advanced prostate cancer, inhibits the NF-kB pathway by inhibiting the IκBα kinase. Decreased DRAIC expression predicted poor patient outcome in gliomas and seven other cancers. We now report that DRAIC suppresses invasion, migration, colony formation and xenograft growth of glioblastoma derived cell lines. DRAIC activates AMPK by downregulating the NF-κB target gene GLUT1, and thus represses mTOR, leading to downstream effects such as decrease in protein translation and increase in autophagy. DRAIC, therefore, has an effect on multiple signal transduction pathways that are important for oncogenesis: the NF-κB pathway and AMPK-mTOR-S6K/ULK1 pathway. The regulation of NF-κB, protein translation and autophagy by the same lncRNA explains the tumor suppressive role of DRAIC in different cancers and reinforces the importance of lncRNAs as emerging regulators of signal transduction pathways.

## Introduction

Glioblastoma (GBM) is a highly invasive, migratory and aggressive form of primary malignant tumor in the central nervous system and responsible for patients morbidity and mortality (1–3). The highly invasive nature of GBM makes these tumors a major challenge for surgical resection. Despite ionizing radiation and chemotherapy agents like Temozolomide, the average survival of GBM patients is only 12-15 months. Therefore, there is an urgent need to better understand the biology of GBMs in order to develop more effective therapies for patients (4–6).

Long noncoding RNAs (lncRNAs) are a class of RNA transcripts that are generally >200 nt in length and do not contain long Open Reading Frames (ORF). Next generation sequencing discovered a vast number of lncRNAs transcribed from different parts of the genome (7,8). LncRNAs regulate gene expression transcriptionally both through modulating the epigenetic state and post-transcriptionally (9–12). Abnormal lncRNA expression is often associated with tumor formation, drug resistance and metastasis (13–15). It is now well established that lncRNAs can act as tumor promoters (oncogenic) or tumor suppressors and regulate tumor progression and development (16–18).

We reported that DRAIC expression is predictive of favorable outcome in prostate cancers, gliomas and six other cancers(18) and that the DRAIC lncRNA exerts a tumor suppressive effect on prostate cancer cells *in vitro* through inhibiting the oncogenic transcription factor NF-κB(17). DRAIC interacted with the IKK complex, specifically with IKKα and NEMO, and disrupted the integrity of the complex, thereby inhibiting IκBα phosphorylation and the downstream NF-κB signaling pathway. DRAIC knockdown/knockout induced NF-κB, cell invasion and soft agar colony formation, and inhibition of NF-κB either by Bay11-7082 or super repressor IκBα suppressed these phenotypes in prostate cancer (17). We have now experimentally tested whether the tumor suppressive effect of DRAIC can be seen *in vitro* on another cancer, GBM, and elucidated the signal transduction pathway by which this action is mediated.

Mammalian target of Rapamycin (mTOR) is a master regulator of cellular protein translation and cell growth(19,20). mTOR is activated by mutations in its regulators in multiple cancers including GBM(21). MTOR is a serine-threonine kinase of the PI3K family that is activated by different external stimuli like insulin, amino acids and different growth factors and controls protein translation. MTOR acts via two distinct complexes, mTORC1 and mTORC2. MTORC1 complex consists of five components, of which we want to highlight regulatory-associated protein of mTOR (Raptor). Raptor is often considered an adaptor protein for recruiting mTOR substrates like eukaryotic initiation factor 4E -binding protein 1 (4E-BP1) and the p-70 ribosomal S6 kinase 1 (S6K1). MTORC1 stimulates protein translation by phosphorylating 4E-BP1 at threonine residues 37 and 46 (Thr37/46), which releases EIF4E for it to associate with mRNA caps and thus increases cap-dependent protein translation. MTORC1 also phosphorylates S6K1 at threonine residue 389 (T389), which promotes protein translation.

mTOR is also known to regulate autophagy, a cellular catabolic process by which cells also maintain energy homeostasis. mTOR inhibits autophagy (22,23) by phosphorylating ULK1 at serine residue 757 (S757), which disrupts the interaction between ULK1-AMPK and prevents the activating phosphorylation on ULK1 serine 555 (S555) by AMPK (24). While inhibiting protein translation usually inhibits tumor progression, the role of autophagy in tumor progression is highly context dependent (25,26), For example, inhibition of autophagy using Bafilomycin A1 and Chloroquine potentiates the effects of chemotherapeutic drugs in triple negative breast cancer (27), and activation of autophagy is a recognized mechanism of chemo-resistance of glioblastomas to temozolomide (28,29). On the other hand, autophagy is also responsible for autophagic apoptosis, a known mode of cancer cell killing after chemotherapy and radiotherapy of GBM. Therefore repression of autophagy may also lead to poor outcome in GBMs (Taylor *et al*., 2018).

AMP-activated protein kinase (AMPK) is a metabolic energy sensor that maintains cellular energy homeostasis in presence of energy stress (30,31). AMPK is a heterotrimeric enzyme that becomes active during energy stress when intracellular concentration of ATP drops and AMP level increases. AMP binds with the γ regulatory subunit of AMPK and allosterically activates the AMPK and accelerates AMPK phosphorylation on threonine 172(Thr172) by the upstream kinase LKB1. Phosphorylation of Thr172 of AMPK enhances its activity several fold further (32). AMPK inactivates mTORC1 which lead to the inhibition of protein synthesis and cell growth and an increase in autophagy (24,33,34). AMPK inhibits mTORC1 by phosphorylating TSC2, an upstream negative regulator of mTORC1 and also by directly phosphorylating Raptor at serine 792 (S792) residue (30,35). As mentioned above, AMPK can also phosphorylate ULK1 at S555 residue to induce the autophagy (36). In addition, AMPK activation by AICAR leads to an increase in FoxO3a phosphorylation at residue serine 413 (S413) and upregulation of autophagy regulated genes, LC3B-II, Gabarapl1 and Beclin 1 (37).

We report that DRAIC is also tumor suppressive on GBM cells *in vitro* and that it regulates another signal transduction pathway in GBM and prostate cancer cells. DRAIC exerts its tumor suppressive function through transmission of the signal from IKK/NF-κB to the AMPK/mTOR pathway via regulation of GLUT1 expression. The inhibition of mTOR by this pathway then leads to inhibition of protein translation and cellular invasion and activation of autophagy.

## Results

### DRAIC overexpression represses cell migration and invasion in glioblastoma cells

Glioblastoma (GBM) cells often show enhanced cell migration and invasion (1,38). Prior to determining the effect of DRAIC on GBM cell migration and invasion, we measured DRAIC expression levels in normal immortalized astrocytes, GBM stem cells and GBM cell lines by qRT-PCR. Endogenous DRAIC expression levels in these cells are very low compared to prostate cancer cells **(Fig. S1)**. We prepared DRAIC overexpressing stable GBM cell lines to examine how DRAIC can be tumor suppressive in these cells. **Figure 1A** shows DRAIC overexpression levels in different GBM cell lines. Overexpression of DRAIC in U87 and A172 GBM cells decreased cell migration compared to cells transfected with Empty Vector (EV) (**Fig. 1B-E**). The decreased cell migration by DRAIC overexpression in A172 and U87 cells was further validated by Time Lapse Video microscopy and shown in Movies (Fig. S1–S4). Similar to the decrease in cell migration, DRAIC overexpression in U87, U373 and A172 cells was also associated with reduced cell invasion through Matrigel in a Boyden Chamber assay **(Fig. 1F-K)**. These results show that DRAIC overexpression in glioblastoma cells decreases cell migration and invasion.

**Figure 1:**
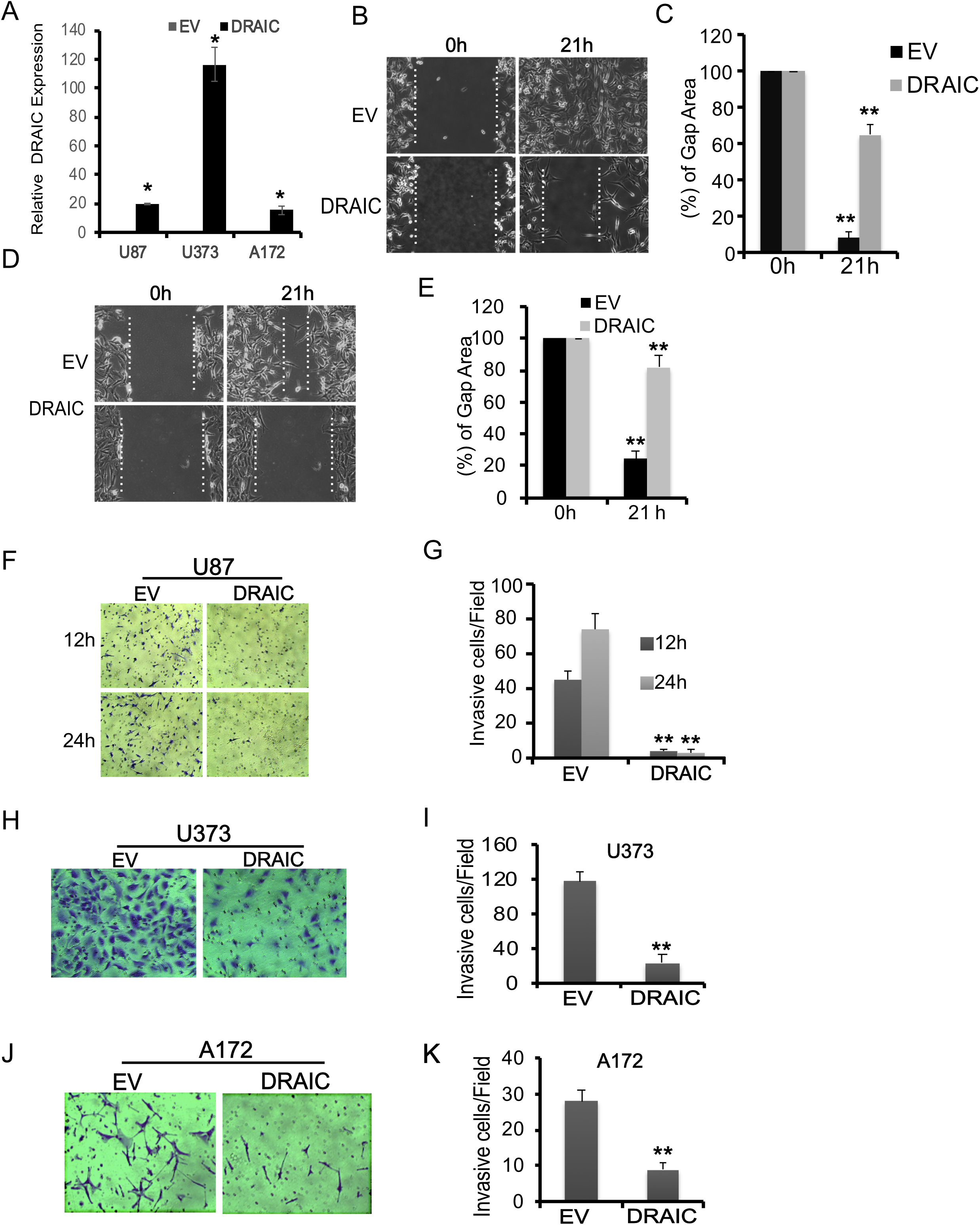
DRAIC suppresses glioblastoma cell migration and invasion: **A-D)** A172 and U87 GBM cells were stably transfected with either empty vector or full length DRAIC. The monolayer cells were scratched using 200 ul filter tips and allowed the cells to migrate over 21 hrs. The images were captured at 0 and 21 hrs time point. The scratch width or gap area was calculated using ImageJ software and quantified in Figure1B and 1D. **E-K)** Matrigel Invasion assay was performed with DRAIC overexpressing U87 **(F-G)**, U373 **(HI)** and A172 **(J-K)** cells. The average number of invasive cells were quantified by counting cells from 10 random fields for each cell lines. Data are presented as mean ± SD *, p<0.05, **, p<0.01.

### DRAIC overexpression suppressed tumorigenicity *in vitro* and *in vivo*

Stable overexpression of DRAIC in U87 and U251 cells did not affect cell proliferation when cells were cultured adherent to a plastic dish **(Fig. 2A, Fig. S2)** but anchorage independent growth was attenuated as evident from a decrease in soft agar colony size and colony number as compared to empty vector expressing cells (EV) **(Fig. 2B-E**). We next examined the effect of DRAIC on tumor growth *in vivo*. U87 cells stably transfected with either EV or DRAIC expressing vectors were stereotactically injected into striata of 6 weeks old immunodeficient mice and tumor growth was followed by MRI. Three weeks after the injection, the EV expressing U87 group showed large tumors, which are marked in the figure with red arrows, with an average tumor volume of 0.65 ± 0.01 mm^3^ **(Fig. 2F, H)**. In contrast, DRAIC overexpressing U87 cells showed a dramatic reduction of tumor volume to 0.02 ± 0.001 mm^3^ **(Fig. 2G, H)**. These results led us to conclude that DRAIC overexpression in GBM cells decreased tumor growth both *in vitro* and *in vivo*.

**Figure 2:**
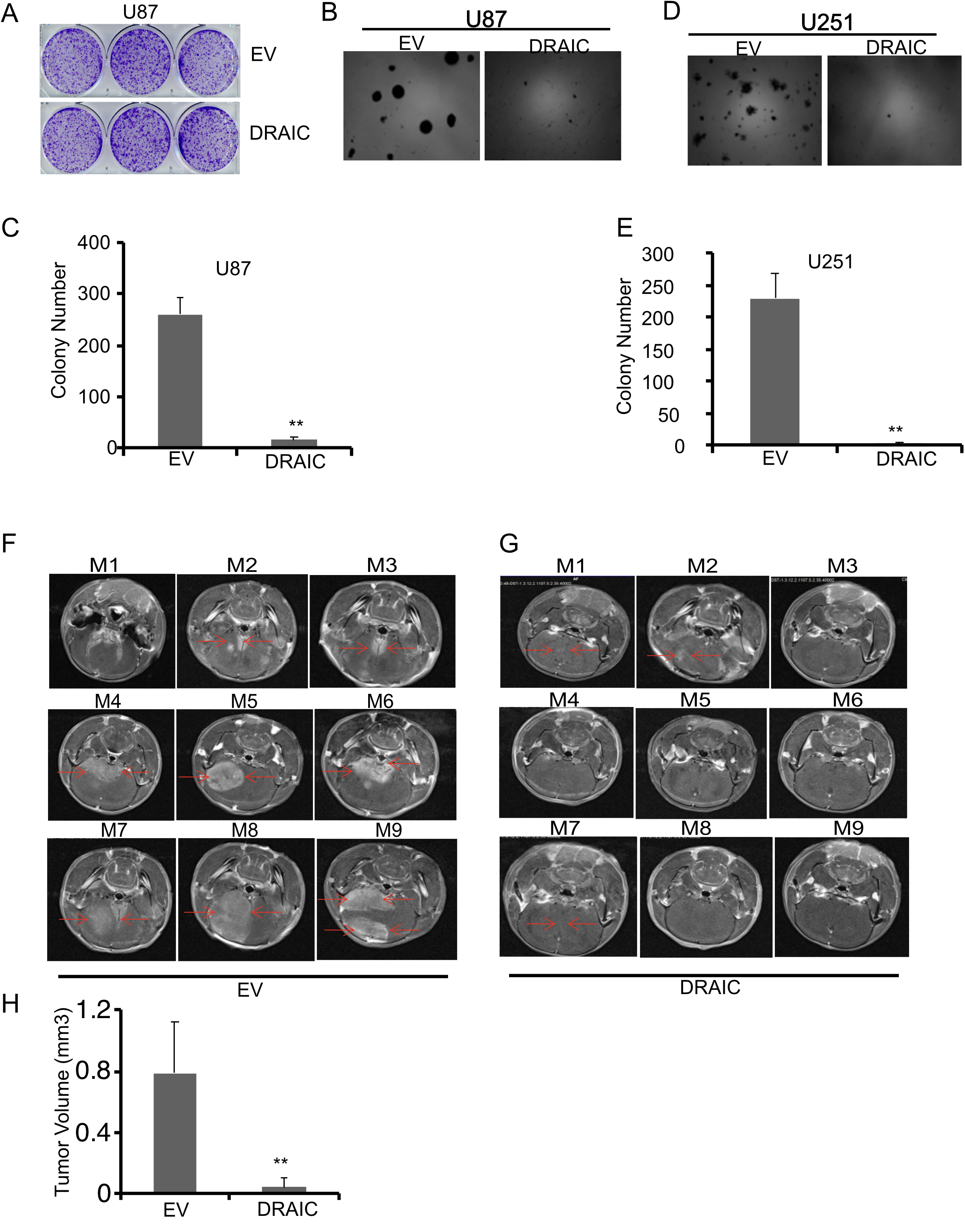
DRAIC overexpression represses the tumorigenic property of glioblastoma cells both *in vitro* and *in vivo*: **A)** Colony formation assay was carried out with EV and DRAIC overexpressing U87 cells. **B-E)** Anchorage-independent soft agar colony formation was executed in U87 (B, C) and U251 cells (D,E). 1X10^4^ cells were seeded for the soft agar assay and monitor the growth of the cells over 3 weeks (B,D). The colony number was quantified by taking an average of 10 random field (C,E). **F, G)** 2 X10^5^ U87 cells either transfected with EV (F) or full length DRAIC (G) were sterostactically implanted into the stratia of immunodeficient mice brain. 3 weeks after the injection, the mice were imaged by MRI scan and quantified in G. Data are presented as mean ± SD *, p<0.05, **, p<0.01.

Note that because of the undetectable levels of DRAIC in GBM cell lines or in GBM derived stem cells, although overexpression of DRAIC can be studied in GBM and prostate cancer cells, knockdown or knockout experiments below have only been done in prostate cancer cells where DRAIC expression is detected (17).

### DRAIC regulates global cellular translation

While screening for various oncogenic properties in the cells, we noted that global cellular translation as measured by puromycin pulse labeling is regulated by DRAIC. Overexpression of DRAIC in both GBM and prostate cancer cells led to decreased global cellular translation **(Fig. 3A, B**). Conversely, knockdown or knockout (KO) of DRAIC in prostate cancer cells stimulated protein translation **(Fig. 3C, D**). Since high mTORC1 activity in tumor cells is often associated with increased cellular translation (39,40), we examined whether mTORC1 is activated when DRAIC is knocked down or knocked out. The phosphorylation of mTORC1 at S2448 residue is sometimes considered a surrogate marker for mTOR activation. DRAIC overexpression did not reduce mTOR phosphorylation at this residue (**Fig. 3E**), but decreased the phosphorylation of mTOR substrate S6K1 (T389) (**Fig. 3E**). Knockout of DRAIC in prostate cancer cells again did not affect the mTOR phosphorylation at S2448 residue but increased the phosphorylation of S6K1 and of S6K substrate S6 **(Fig. 3F**). Consistent with regulation of mTORC1, the phosphorylation of another substrate mTORC1 substrate, ULK1 on S757, was decreased by overexpression of DRAIC and increased by knockout of DRAIC (**Fig. 3G, H**) These results suggest that DRAIC regulates cellular phenotypes, including global protein translation, by modulating the activity of mTOR kinase.

**Figure 3:**
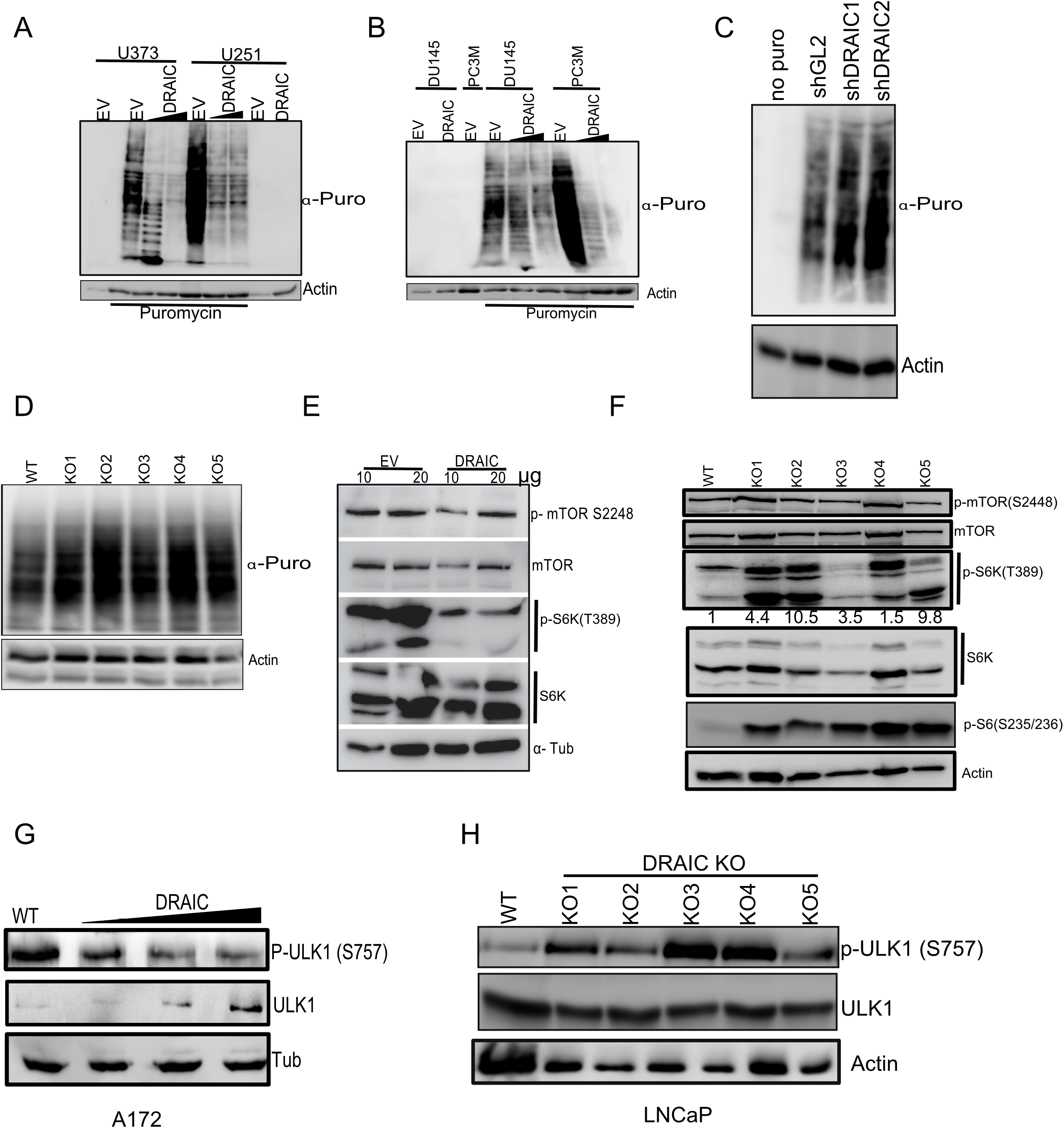
Low DRAIC expression is associated with increased cellular translation in glioblastoma: **A,B)** U373, U251 (A), DU145 and PC3M cells (B) were transfected with either EV or DRAIC followed by 10 ug/ml puromycin pulse labeling for 1 hr and immunoblotted with antibodies against puromycin and loading control actin. **C, D)** LNCaP either stable DRAIC knockdown (C) or knockout (D) of DRAIC cells were pulse labeled with puromycin for 1 hr and cell lysates were subjected to immunoblot with the antibodies against puromycin and actin. **E)** U87 cells were transfected with either EV or full length DRAIC and 10 and 20 *μ*g of cell lysates was loaded and immunoblotted with antibodies against phospho mTORC1 (S2448), phospho S6K (T389), total S6K and alpha tubulin. **F)** EV and DRAIC KO LNCaP cell lysates were immunoblotted with antibodies against phospho mTORC1 (S2448), phospho S6K (T389), total S6K, phospho S6 (S222/S223) and loading control actin. G) A172 cells were stably transfected with EV and DRAIC overexpressing plasmids and cell lysates were immunoblotted with antibodies phospho ULK1 S757, ULK1 and tubulin. H) Different KO single clones of DRAIC were immunoblotted with antibodies against phospho ULK1, ULK1 and actin.

### DRAIC regulates AMPK kinase and thus regulates mTORC1

The mTOR pathway is regulated indirectly by several upstream kinases (41), particularly by AMPK (42,43). The schematic shows the different substrates which are phosphorylated and regulated by AMPK **(Fig. 4A**). To understand the mechanism of decreased mTOR substrate phosphorylation upon DRAIC overexpression, we checked AMPK phosphorylation status at T172 residue, a marker for AMPK kinase activity. Overexpression of DRAIC in A172 GBM cells increased AMPK phosphorylation (**Fig. 4B**) whereas DRAIC KO in LNCaP cells decreased AMPK phosphorylation **(Fig. 4C**). One of the mechanisms by which AMPK inhibits mTOR activity is by phosphorylating Raptor at S792 residue (30). Overexpression of DRAIC in A172 and U251 cells increased Raptor phosphorylation on S792 (**Fig. 4D, G**) whereas DRAIC KO decreased Raptor phosphorylation on S792 **(Fig. 4C**). Two other substrates of AMPK, ULK1 (S555), and FoxO3A (S413), were phosphorylated upon DRAIC overexpression suggesting that AMPK activity is globally induced upon DRAIC overexpression **(Fig. 4E-I)** while DRAIC KO decreased level of phospho FoxO3A S413. Together these results suggest that DRAIC stimulates AMPK, and this inhibits mTORC1 to inhibit phosphorylation of key substrates like S6K1, which inhibits translation.

**Figure 4:**
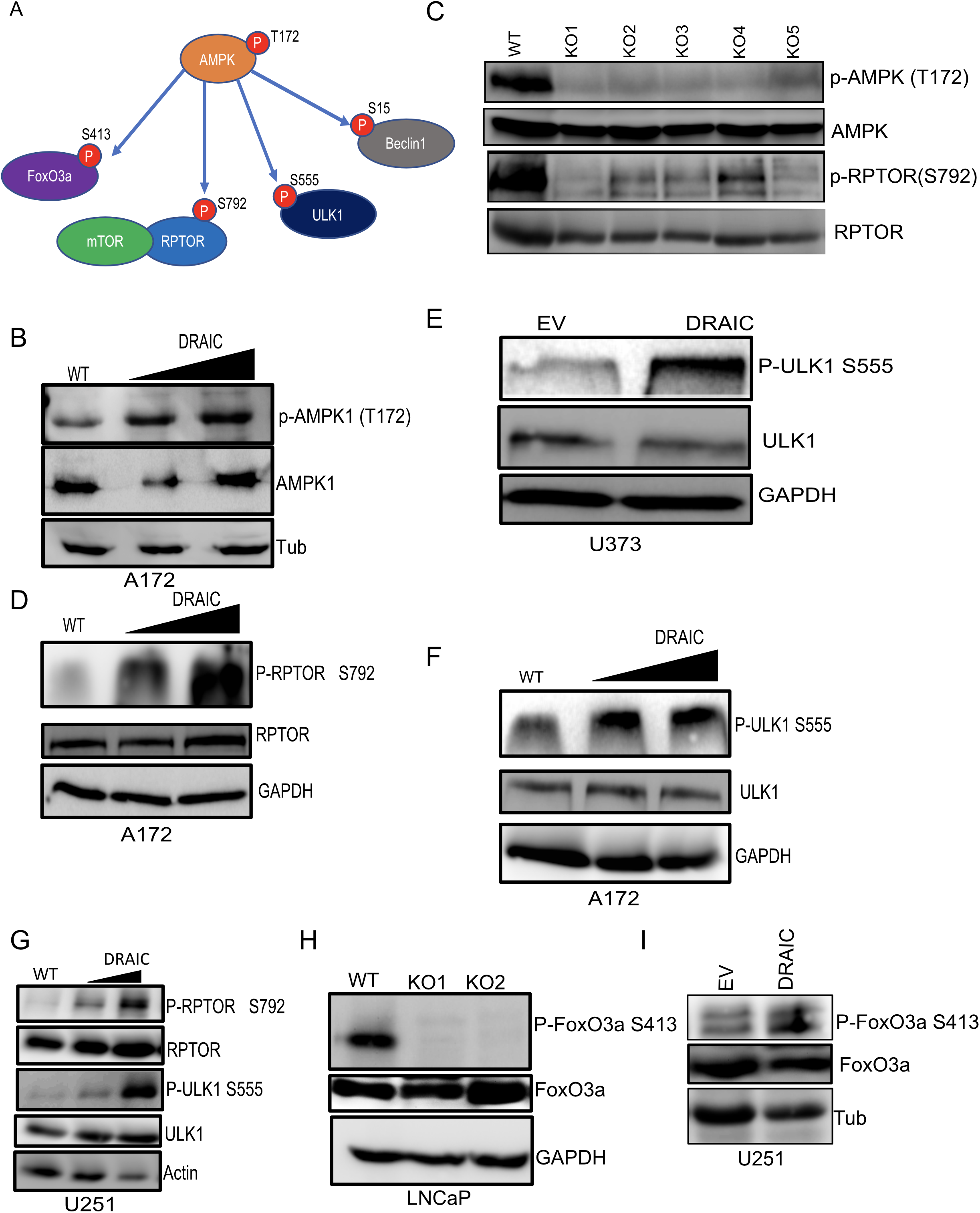
DRAIC regulates AMPK substrate phosphorylation: **A)** Schematic representation of different substrates, which were phosphorylated by AMPK. **B)** Empty vector (EV) and DRAIC overexpressing A172 cells were immunoblotted with antibodies against phospho AMPK (T172), total AMPK and loading control tubulin. **C)** WT and multiple DRAIC KO clones were subjected to immunoblot with antibodies against phospho AMPK (T172), total AMPK, phospho Raptor (S792) and total Raptor. **D, E)** EV or DRAIC overexpressing U373 cells were immunoblotted with antibodies against phospho Raptor (S792), total Raptor, GAPDH (D) and (E) phospho ULK1 (S555), total ULK1 and loading control GAPDH. **F)** EV and stable DRAIC overexpressing A172 cells were subjected to immunoblot with antibodies against phospho ULK1 (S555), total ULK1 and loading control GAPDH. **G)** U251 cells stably transfected DRAIC was immunoblotted with antibodies against phospho RPTOR S792, RPTOR, phospho ULK1 S555, ULK1 and loading control acting. **H, I)** DRAIC KO LNCaP cells and U251 cells stably transfected with EV or DRAIC were subjected to western blotting with phospho FoxO3a (S413).

### DRAIC overexpression induces autophagy and associated gene expression

AMP activated protein kinase (AMPK) regulates cellular energy metabolism and homeostasis by controlling autophagy(33,34). Indeed, the increase in ULK1 S555 and FoxO3A S413 phosphorylation upon DRAIC overexpression (**Fig. 4**) should promote autophagy. DRAIC overexpression in U251 cells decreased the levels of LC3 II and P62 the core proteins involved in autophagosome formation, which are subsequently degraded when the autophagosomes fuse to lysosomes (**Fig. 5A)**. To distinguish whether the decrease of LC3 II level is due to the inhibition of autophagosome formation or increased lysosomal degradation, DRAIC overexpressing cells were treated with bafilomycin A1 to inhibit lysosomal degradation. Bafilomycin A1 restored the level of LC3 II in DRAIC overexpressing cells, suggesting that lysosomal degradation of LC3II and by extension, autophagic flux, is increased when DRAIC is elevated in GBM cells (**Fig. 5B lane 1 with 2 and 3)**. Knockout of DRAIC in LNCaP cells increased LC3 II expression suggesting the inhibition of autophagy (and inhibition of lysosomal degradation) of LC3 II. Consistent with a decrease in autophagic flux upon DRAIC decrease, the level of P62 is also increased in the DRAIC KO prostate cancer cells (**Fig. 5C**).

**Figure 5:**
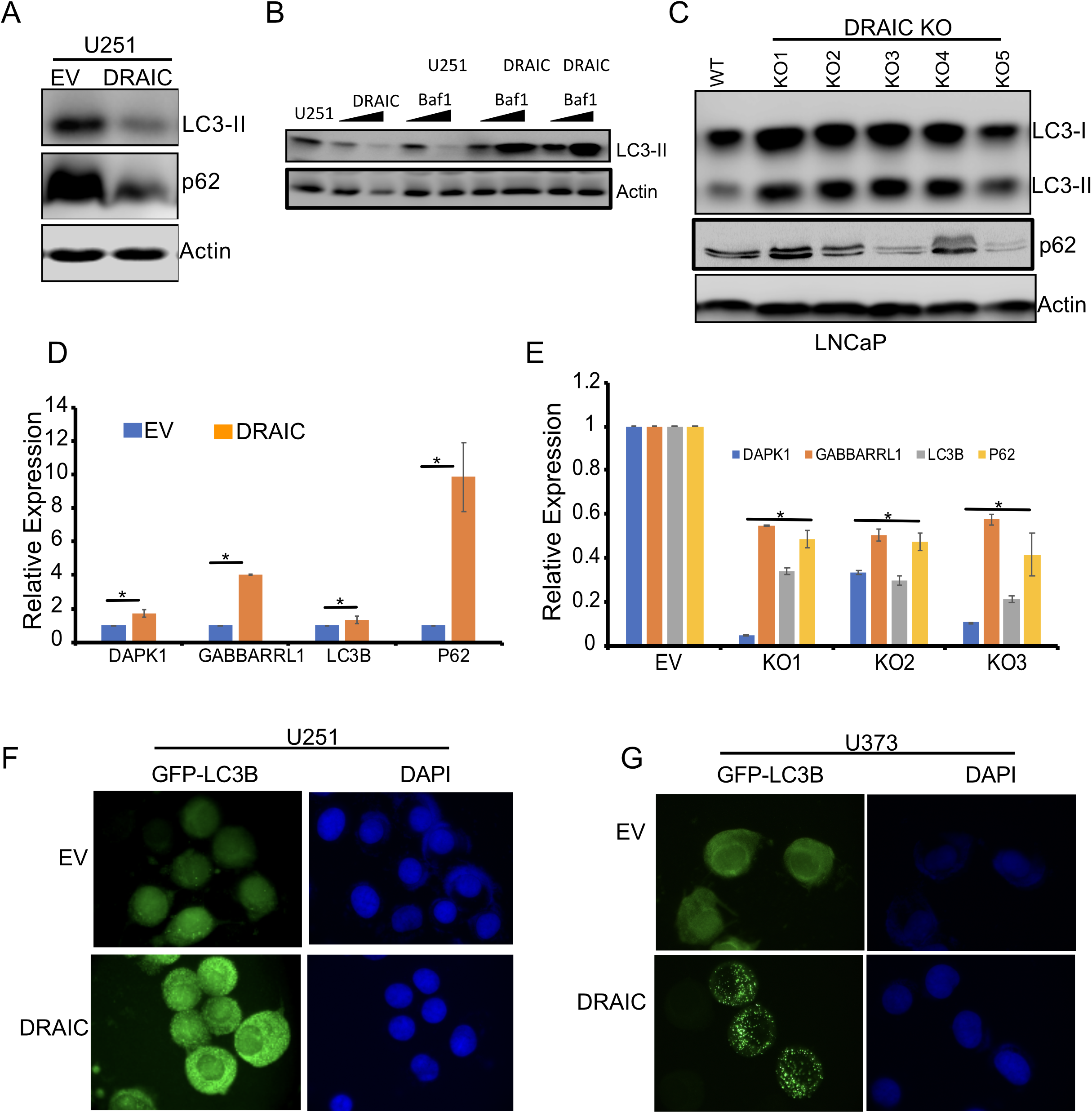
Overexpression of DRAIC induces the autophagy: **A)** U251 cells transfected with EV and DDRAIC were immunoblotted with antibodies against LC3B, p62 and internal loading control actin. **B)** U251 cells were stably transfected with empty vector (EV) or full length DRAIC were treated with 10 nM of bafilomycin A1 for 24 hrs and cell lysates were subjected to immunoblot with LC3B and actin antibodies. **C)** WT and DRAIC KO LNCaP clones were immunoblotted with antibodies against LC3B, p62 and actin. **D, E)** qRT-PCR analysis of autophagy responsive gene expression in U251 cells (D) and DRAIC KO LNCaP cells (E). **F,G)** DRAIC overexpressing U251 (F) and U373 (G) cells were transfected with GFP tag LC3B and puncta formation was assessed to monitor the autophagosome formation. Data are presented as mean ± SD *, p<0.05.

The DRAIC-AMPK pathway increased the phosphorylation and activation of FoxO3A, a transcription factor important for induction of genes associated with autophagy (**Fig. 4H, I**). Consistent with this, DRAIC overexpression increased the expression of autophagy-associated genes, whereas DRAIC KO decreased expression of those same genes (**Fig. 5D-E**,).

Finally, we performed microscopic imaging of LC3 by fusing it with a GFP tag to express a chimeric GFP-LC3B that can mark autophagosome formation as fluorescent puncta. DRAIC overexpression in both U251 and U373 cells increased GFP-LC3B puncta consistent with an induction of autophagy **(Fig. 5F-G)**.

Overall, these results suggest that DRAIC overexpression leads to an increase in autophagy while decreasing protein translation, and conversely a decrease in DRAIC, as in the DRAIC KO cells, decreases autophagy along with an increase in translation. Thus DRAIC elevation mimics conditions of energy stress or amino-acid starvation, and this may be responsible for the suppression of oncogenic phenotypes.

### DRAIC KO induced protein translation and cellular invasion are reversed by activating AMPK

To understand whether increased protein translation seen upon DRAIC KO LNCaP is mediated by decrease in active AMPK, we pre-treated both WT and DRAIC KO LNCaP cells with AMPK activators AICAR or metformin for 24 hr and performed the global protein translation assay with puromycin pulse labeling. Both AMPK activators rescued the DRAIC KO mediated increase in global protein translation (**Fig. 6A**). The WT and DRAIC KO LNCaP cells were also treated with different concentrations of rapamycin, a well-known inhibitor of mTORC1 and cellular translation, as a positive control in our experiment (**Fig. 6A**).

**Figure 6:**
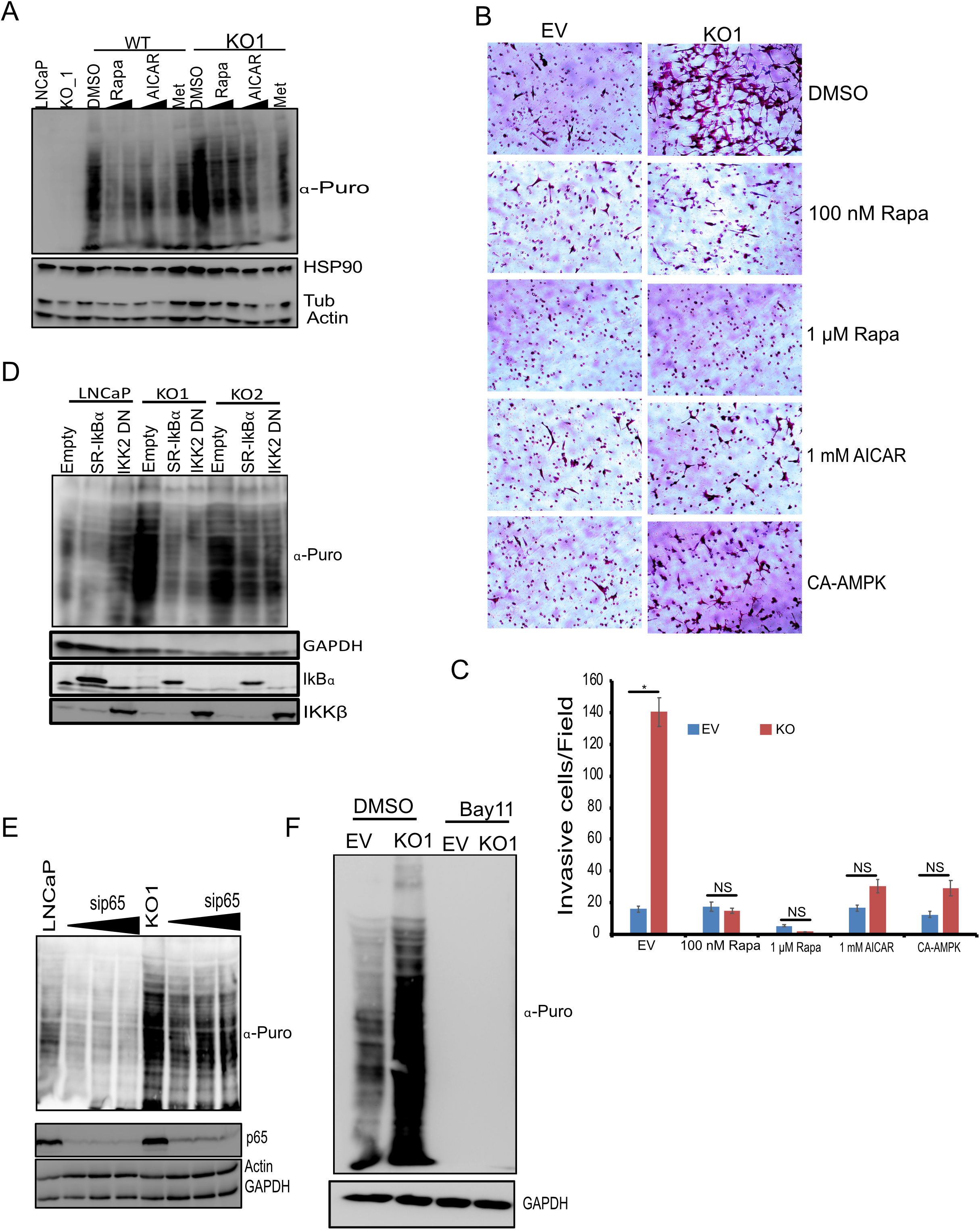
DRAIC KO induced translation and invasion were rescued by AMPK activators: **A)** LNCaP and DRAIC KO cells were pre-treated either with translation inhibitor, rapamycin or 1 mM AMPK activators AICAR and metformin for 24 hrs followed by puromycin pulse labeling and immunoblotted with puromycin antibody. **B, C)** WT and DRAIC KO LNCaP cells transfected with constitutive active AMPK or pre-treated with 0.1, 1 μM rapamycin or AMPK activator AICAR for 24 hrs and Boyden chamber invasion assay was performed. 10 random fields were selected for counting the number of invasive cells per field. **D)** LNCaP and DRAIC KO cells were transfected with either super repressor IκBα (SR-IκBα) or IKK beta dead kinase mutant (IKK2-DN) for 48 hrs followed by puromycin pulse labelling and immunoblotted with puromycin antibody. **E, F)** LNCaP and DRAIC KO cells were transfected with siRNA against p65 or treated with IKK inhibitor Bay11-7082 for 2 hours followed by puromycin pulse labeling and immunoblotted with antibodies against puromycin, p65, actin and GAPDH. Data are presented as mean ± SD *, p<0.05.

We also checked whether increased cellular invasion seen upon DRAIC KO is because of AMPK inhibition, by inducing AMPK activity by AICAR or by overexpressing constitutive active AMPK, and then performing the Matrigel Boyden chamber invasion assay. Both the chemical or genetic activation of AMPK led to a decrease in cell invasion in DRAIC KO LNCaP cells (**Fig. 6B, C Fig. S3**). Here again, rapamycin treatment demonstrates that the increase in protein translation (downstream of AMPK inhibition) also decreases the cellular invasion in DRAIC KO LNCaP cells.

Taken together, these results suggest that AMPK inhibition in DRAIC-deficient cells is responsible for the increase in cellular translation and increase in invasion (**Fig. 6B, C, Fig. S3)**. The effects of rapamycin on protein translation and invasion suggest that the increased invasion is dependent on the increase in protein translation.

### The interplay of IKK-NF-kB with AMPK-mTOR-protein translation and cell invasion

DRAIC KO, by inhibiting IKK activates the NF-κB pathway (17). We previously showed that DRAIC KO induced invasion could be rescued by inhibiting IKK and NF-κB. Above, we showed that the increase in invasion and protein translation can also be rescued by activating AMPK. This raises the question whether IKK and the NF-κB pathway is involved in transmitting the signal from DRAIC through AMPK to protein translation/cellular invasion. To understand the involvement of DRAIC in attenuating cellular translation through NF-κB pathway, we inhibited NF-κB in DRAIC KO cells by four strategies: transfecting plasmids overexpressing IκBα super repressor (S32A/S36A) or the catalytically dead IKKβ kinase (IKK2-Dominant Negative), knocking down NF-κB P65 by siRNA or treated with IKK inhibitor, Bay11-7082. Inhibition of NF-κB pathway by any of the above strategies reversed the increase in protein translation in DRAIC KO cells (**Fig. 6D-F**).

### Increase in cellular invasion and translation in DRAIC KO LNCaP cells is reversed by inhibiting NF-kB target gene GLUT1

We next asked how NF-κB activity (seen upon DRAIC KO) could suppress AMPK activity. Increase in NF-kB target gene expression is often associated with increased tumorigenicity and metastasis (44). NF-kB and IKK are known to increase the expression and surface localization of the NF-κB target gene GLUT1 (45), which would result in increased glucose import and so a decrease in the AMP:ATP ratio in cells, which may be sufficient to decrease AMPK activation. To explore the involvement of GLUT1 in regulating AMPK activity, we assessed the expression of GLUT1 in DRAIC KO and overexpressing cells. GLUT1 expression is increased in DRAIC KO cells and deceased in DRAIC overexpressing cells (**Fig. 7A, Fig. S4**). The increase in cellular invasion and translation seen in DRAIC KO cells can also be reversed by inhibiting GLUT1 with the pharmacological inhibitor Bay-876 (**Fig. 7B-D**). We conclude that increased GLUT1 expression is responsible for induction of cellular invasion and induction of protein translation in DRAIC KO cells.

**Figure 7:**
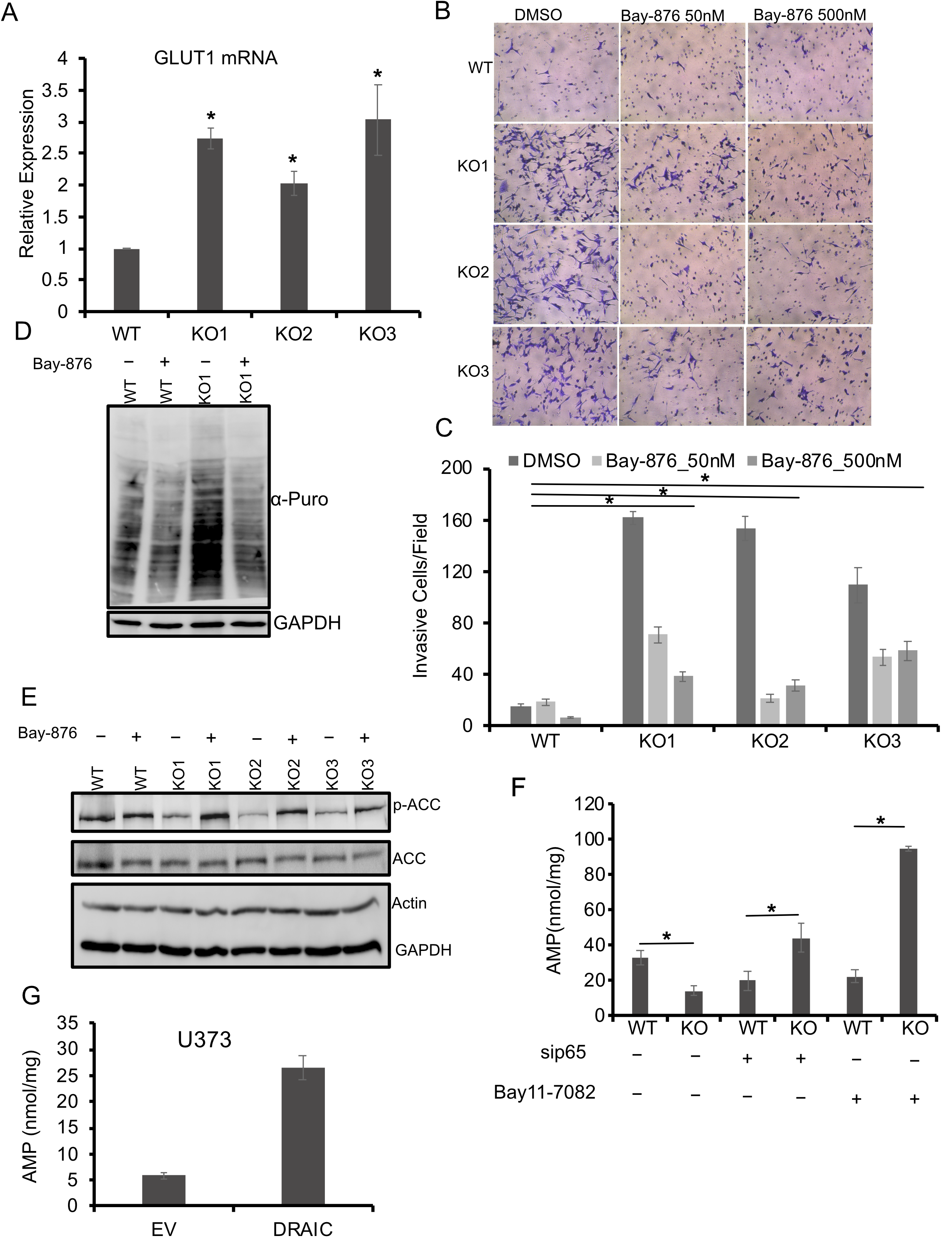
Induction of protein translation, invasion and AMP target phosphorylation in DRAIC KO cells is reversed with GLUT1 inhibitor: **A)** GLUT1 mRNA expression in DRAIC KO LNCaP cells were measured by RT-qPCR. **B, C)** WT and DRAIC KO clones were treated with GLUT1 inhibitor, Bay-876 for 24 hours and Boyden Chamber Matrigel invasion assay was carried out. The invasive cells were strained with crystal violet and Images were captured under Microscope. 8 random field was captured and quantified. **D)** DRAIC KO cells were treated with GLUT1 inhibitor, Bay-876 for 24 hours followed by pulse labeling with puromycin and immunoblotted with antibody against puromycin. **E)** WT and DRAIC KO clones were treated with Bay-876 for 24 hours and immunoblotted with antibodies against phospho ACC S79, total ACC and internal loading control GAPDH. **F)** 2X10^7^ cells were lysed with AMP lysis buffer and AMP concentration was measured using BioVision kit. **G)** AMP concentration in EV and DRAIC overexpressing U373 cells according to manufacturer protocol. Data are presented as mean ± SD *, p<0.05.

As DRAIC KO decreased AMPK phosphorylation at T172 residue, we asked whether this effect is due to the increase in GLUT1 expression, which is known to regulate the activity of AMPK by regulating the ratio of AMP/ATP. Increased AMP/ATP ratio leads to allosteric activation of AMPK through the binding of AMP to the γ subunit of AMPK. To assess whether the elevation of GLUT1 is responsible for the inhibition of AMPK activity we followed AMPK activity by immunoblotting for another AMPK target, phospho ACC Ser 79. DRAIC KO cells show decreased phospho ACC Ser 79 expression, consistent with the low activation of AMPK. However, GLUT1 inhibitor treatment of the DRAIC KO cells rescued the ACC phosphorylation, suggesting that the inhibition of AMPK was indeed mediated by the increase of GLUT1 (**Fig. 7E**). To add to the evidence, we measured the cellular AMP level in DRAIC KO cells and discovered that the AMP concentration is decreased in DRAIC KO cells and that this reduction in AMP is rescued by inhibiting the NF-κB pathway either with IKK inhibitor Bay11-7082 or knocking down p65 (**Fig. 7F**). In contrast, overexpression of DRAIC in U373 cells, which decreases GLUT1 expression (**Fig. S4**), leads to an increase of AMP level **(Fig. 7G**). These results suggest that increased GLUT1 expression in DRAIC KO cells leads to reduction of AMP/ATP ratio, which decreases AMPK activity and thus increases cellular translation.

## Discussion

Our results identify a major signal transduction pathway through which DRAIC mediates its tumor suppressive effect on cancer cells of different lineages. As in prostate cancer, DRAIC suppresses glioblastoma development both *in vitro* and *in vivo*. We discovered the following: 1. Overexpression of DRAIC in different glioblastoma cells decreased cell migration, cell invasion, anchorage-independent growth and xenograft growth. 2. DRAIC decreases protein translation through inhibiting the phosphorylation of mTORC1 substrates. 3. DRAIC induces AMPK phosphorylation at T172 residue, which is known to repress mTORC1 to repress translation and phosphorylates several substrates that lead to an increase in autophagic flux. 4. The increase in translation after DRAIC knockout is reversed by either inhibiting IKK kinase activity, knocking down p65 or activating AMPK. Taken together these results support a model (**Fig. 8**) where DRAIC inhibits several tumor phenotypes in prostate cancers and gliomas by inhibiting NF-κB, and thus activating AMPK, inhibiting mTORC1, inhibiting protein translation and increasing autophagy. The link between NF-κB and AMPK is significantly mediated by the ability of NF-kB to induce the glucose transporter GLUT1. Thus, when DRAIC is decreased in tumors, NF-kB is activated, leading to increased glucose uptake and decreased AMP/ATP ratio and repression of AMPK. This results in increase of protein translation through mTORC1 activation and inhibition of autophagy through AMPK repression. So, at least in this context, inhibition of autophagy appears to make tumors more aggressive, consistent with the notion that autophagic apoptosis is important for GBM cell death following temozolamide and radiotherapy.

**Figure 8:**
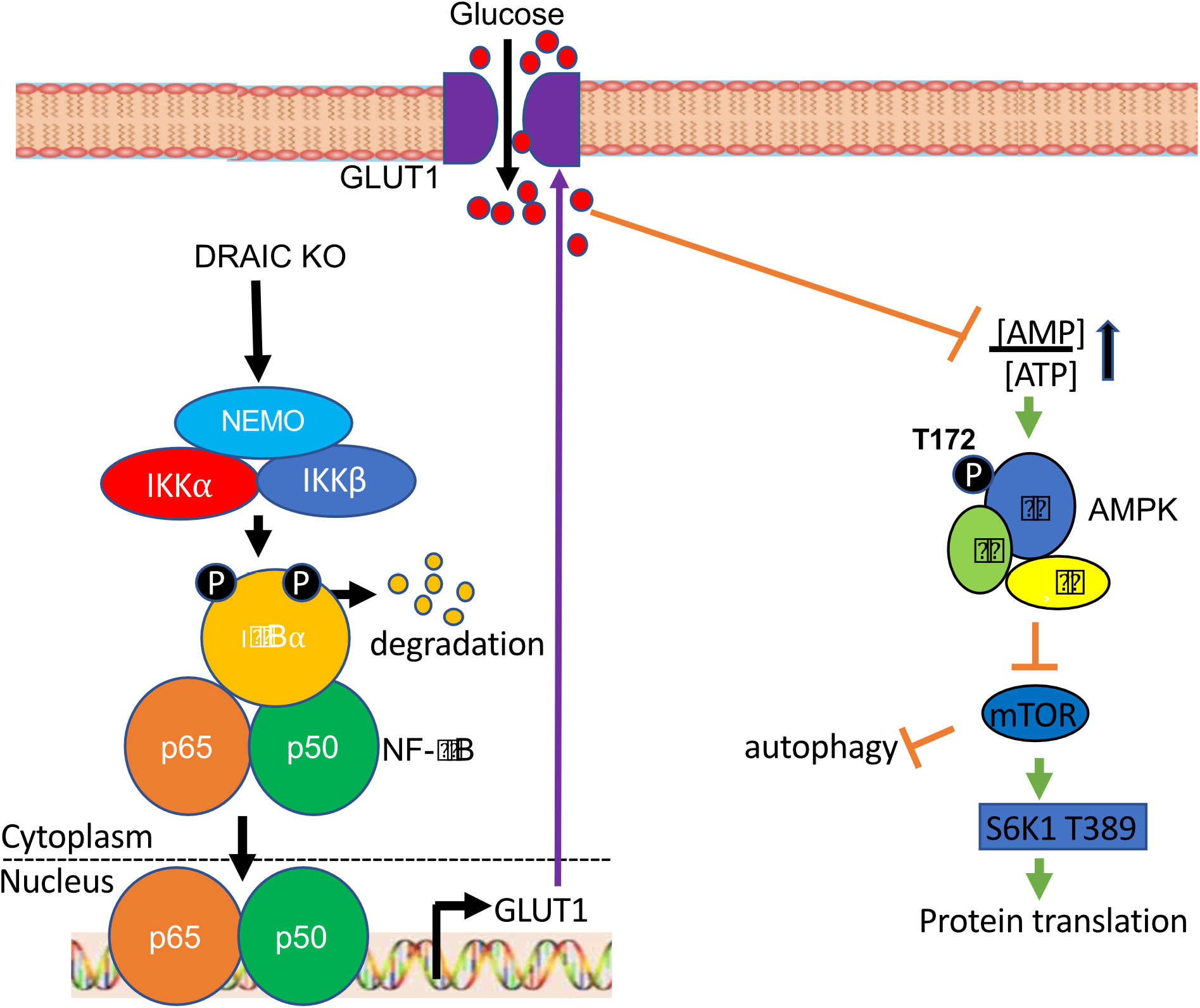
Model of the pathway by which DRAIC regulates translation, autophagy, migration and invasion and thus acts as a tumor suppressor.

It is interesting that although previous literature reported that mTOR and Raptor both are associated with the IKK complex (46), our results with AICAR and metformin (activators of AMPK) suggest that the effect of IKK on mTORC1 is mediated through AMPK.

Our results add DRAIC to the few lncRNAs which are known to be involved in regulating the mTORC1 pathway. H19 long noncoding RNA controls pituitary tumor growth by inhibiting mTORC1 pathway. H19 interacts with the TOS domain of 4E-BP1 and masks the Raptor binding site and therefore inhibits the 4E-BP1 phosphorylation by mTORC1 without affecting S6K1 phosphorylation(47). The lncRNA MALAT1 is oncogenic in hepatocarcinomas by inducing the expression of an oncogenic splicing factor SRSF1 which promotes the expression of a short isoform of S6K1 that binds with mTORC1 and activates the kinase (48). HAGLROS directly binds to and activates mTOR and acts as a competing endogenous RNA to antagonize miR-100-5p and thus increases mTOR expression (49). DLEU1 has been shown to associate with mTOR, though it is not clear whether the binding regulates mTORC1(50). In contrast to the other lncRNAs, however, DRAIC does not physically associate with mTOR (not shown) and appears to act on mTOR by regulating its upstream activator AMPK.

Recent literature suggests that there are a few lncRNAs that modulate AMPK activity. The lncRNA NBR2 interacts with AMPK during energy deprivation to activate AMPK, and its expression is induced by LKB1-AMPK thereby producing a feed-forward loop to activate AMPK(51). The oncogenic Lnc-THOR depletion inhibits glioma cell survival by decreasing MAGEA6 mRNA and protein, which leads to decreased degradation of AMPKα by MAGEA6-TRIM28 E3 ubiquitin ligase (52). The increase of AMPK level and activity is detrimental to glioma cell survival. We now report that DRAIC also promotes AMPK activation and suppresses tumor progression. However, here again, we did not see any direct association of DRAIC with AMPK or with any of the activating kinases of AMPK.

We show that DRAIC regulates the AMPK pathway through NF-κB. DRAIC reduces the expression of glucose transporter GLUT1, an NF-κB responsive gene, which leads to decreased uptake of glucose through GLUT1 and increase in the ratio of AMP/ATP. This increased intra-cellular concentration of AMP leads to activation of AMPK pathway and stimulation of autophagy in cells overexpressing DRAIC.

A consistent tumor suppressive role of DRAIC *in vitro* on prostate cancer and glioblastoma suggests that this is an effect generalizable to multiple lineages. This is consistent with the finding that decreased DRAIC expression predicts poor outcome in eight different malignant tumors (18). However, there are a few studies on DRAIC specifically in breast cancer where it may act as an oncogenic lncRNA by regulating different cellular functions but the molecular mechanism behind this oncogenic nature is not clearly understood (53–55). DRAIC knockdown in MCF7 breast cancer cells increased autophagic flux (54). In contrast we note that in both GBM and prostate cancer cells DRAIC overexpression increases autophagic flux and DRAIC knockout increases p62, consistent with a decrease in autophagic flux (**Fig. 5**). The opposite effects that DRAIC appears to have on tumor progression in these lineages (oncogenic in breast, tumor suppressive in prostate and gliomas) could be explained by the lineage-specific opposite effects of DRAIC on autophagic flux. Despite these tissue specific differences, our data suggests that activation of DRAIC expression in prostate cancer and gliomas could significantly improve outcomes in these patient populations. In addition, drugs that activate AMPK (like metformin and AICAR) or inhibit mTORC1 (like rapamycin) should be explored for the therapy of GBMs and other tumors with low (or undetectable) levels of DRAIC.

## Acknowledgement

**T**his work was supported by R01 AR067712 and R01 CA060499 to AD. P30 CA44579 to SS. We acknowledge the MAPS core facility at University of Virginia for helping with the animal experiment. Manikarna Dinda for preparing the graphical summary and for reading the manuscript.

## Author Contributions

**Conception and design**: S. Saha, A. Dutta. **Development of methodology**: S. Saha, A. Dutta. **Acquisition of data** (provided animals, acquired and managed patients, provided facilities, etc.): S. Saha, Y. Zhang, R. Abounader, A. Dutta. **Analysis and interpretation of data** (e.g., statistical analysis, biostatistics): S. Saha, B. Wilson, A. Dutta. **Writing, and editing the manuscript:** S. Saha, A. Dutta. **Visualization and critical reading**: S. Saha, Y. Zhang, B. Wilson, R. Abounader, A. Dutta. **Administrative, technical, or material support** (i.e., reporting or organizing data, constructing databases): S. Saha, A. Dutta. **Study supervision**: A. Dutta.

## Materials and Methods

### Cell culture, transfection and Microscopy

LNCaP, PC3M, DU145, and U373 cells were maintained in RPMI medium containing 10% FBS, 1% penicillin/streptomycin, 1 mmol/L sodium pyruvate, and 10 mmol/L HEPES buffer. HeLa, A172 and U251 cells were grown in high glucose DMEM medium supplemented with 10% FBS and 1% penicillin/streptomycin. U87 cells were cultured in MEM medium with 10% FBS, 1% penicillin/streptomycin, 1 mmol/L sodium pyruvate, 10 ml of 7.5% sodium bicarbonate and 5 ml nonessential amino acids. All the cell lines were procured from ATCC and maintained in humidified incubator at 37°C in the presence of 5% CO_2_. All the cell lines were authenticated by short tandem repeats analysis at 15 genomic loci and the amelogenin gene (Biosynthesis). Stable cell lines expressing full length DRAIC was generated in PC3M, A172, U87, U373, U251 and HeLa cells. 1 μg of EGFP-LC3B plasmid was transfected with Lipofectamine 3000 into U373 and U251 cells overexpressing either EV or DRAIC. After 48 hrs of post transfection, cells were imaged with 63X objective using the Zeiss Fluorescence microscope.

### Time Lapse Video Microscopy

The bright field time lapse video microscopy was performed to monitor the cell migration with Zeiss Observer Z1 wide-field microscope. The live cell movement was captured every 10 min for a duration of 24 hrs using the Zeiss software. The cells were maintained at 37°C in presence of 5% CO2.

### Plasmids and reagents

The plasmids pcDNA3-Flag-LKB1 (item # 8591), pEBG-AMPKα1(1-312) (item #27632) IKK2 K44M ((item #1104), pEGFP-LC3 (item # 24920) were procured from Addgene. Antibodies to IKKα (#ab109749), IKKβ (#ab32135), Ribosomal S6 (#ab 40820), STAT1 (#ab109320) were purchased from Abcam. phospho AMPKα (T172) (#2531), AMPKα (#5831), phospho mTOR (S2448) (#2971), mTOR (#2972), LC3B (#3868), phospho ULK1 (S757) (Cell Signaling, #14202), ULK1 (Cell Signaling, #8054), Phospho Raptor (S792) (#2083), Raptor (#2280), LKB1 (#3050), phospho p70 S6K (T389) (#9234), p70-S6K (#2708), phospho S6 (S235/236), phospho Beclin-1 (S15) (#84996), Beclin-1(#3495), phospho FoxO3a(S413) (#8174), p62 (#23214), phospho Acetryl-CoA Carboxylase Ser79 (#3661), Acetyl-CoA-Carboxylase (#3676) were purchased from Cell Signaling Technology. Raptor (#A300-553A) antibody was purchased from Bethyl Laboratories. Antibody against α-tubulin (#sc-5286), β-actin (#sc-47778), HSP90 (#sc-13119), GAPDH (#sc-47724) were purchased from Santa Cruz Biotechnology. Puromycin antibody (#MABE343) was purchased from Sigma-Millipore. The shRNAs oligos against DRAIC were annealed and cloned into pLSP lentivirus vector. Bay11-7082 (#S2913), AICAR (5-aminoimidazole-4 carboxamide-1-β-D-ribofuranoside) (# S1802), Metformin (#S1950), Rapamycin (#S1039), Bafilomycin A1 (#S1413) were purchased from Selleckchem. Noble agar (#S5431) was purchased from Simga.

### Electrophoresis and Immunoblot analysis

The cells on petri dish were washed twice with cold phosphate-buffered saline (pH 7.4) and lysed with the lysis buffer (50 mM Tris-HCl, pH 8.0, NaCl, 10 mM NaF, 1 mM EDTA (pH 8.0), 0.5% NP-40, 1 mM PMSF, 1 mM DTT, 0.5% sodium deoxycholate, Halt protease inhibitor cocktail (Fisher Scientific, #78429) and phosphatase inhibitor tablet (Sigma, #4906845001). The lysates were centrifuged at 15,1000g for 20 min at 4°C and the supernatant was boiled with 1X Laemmli sample buffer and the different amounts of proteins were separated on 10-12% SDS-PAGE and immunoblotted with antibodies against phospho mTOR S2448 (1:1000), mTOR (1:1000), phospho S6K T389 (1:1000), S6K (1:2000), α-tubulin (1:5000), phospho S6 S235/236 (1:2000), S6 (1:4000), actin (1:5000), puromycin (1:5000), phospho AMPK T172 (1:1000), AMPK (1:2000), phospho Raptor S792 (1:1000), Raptor (1:2000), phospho ULK1 S555(1:1000), ULK1 (1:2000), GAPDH (1:5000), phospho Beclin-1 S15 (1:1000), Beclin-1 (1:2000), phospho p53 (1:1000), p53 (1:4000), phospho FoxO3a S413 (1:1000), LC3B (1:1000), p62 (1:2000), HSP90 (1:5000).

### Colony formation, cell proliferation and invasion assay

U87 and A172 cells overexpressing either EV or DRAIC were used for colony formation assay. Briefly, 1000 cells from each conditions were plated in each well of 6 well plates and incubated at 37°C with 5% CO_2_ incubator for around 2 weeks and colonies were stained with 0.05% crystal violet containing 1% formaldehyde and gently rinse with water. The plates were dried and imaged with a scanner. The MTT assay was performed as described previously (17). Briefly, 5 × 10^4^ cells stably transfected with EV and DRAIC were seeded in 24-well plates and MTT assay was performed at different days. The Matrigel-containing Boyden chamber invasion assay was carried out as described previously (17). EV and DRAIC KO cells were transfected with constitutive active AMPK or pre-treated the cells with rapamycin and AMPK activator AICAR for 24 hrs. The cells were trypsinized and a total of 1 × 10^5^ cells were added in serum-free medium in the top of the chamber and full growth medium containing 10% FBS was added to the bottom of the chamber as a chemoattractant and the chamber incubated at 37°C in presence of 5% CO_2_ for 24 hours. After 16-24 hours, the invaded cells on the bottom surface of the membrane were gently washed with 1× PBS and fixed with 100% methanol for 5 minutes followed by 0.5% crystal violet staining at room temperature for 15 minutes. The noninvading cells from the top surface of the chamber were removed by cotton swab. 10 random fields were captured under microscope and the invaded cell number were quantified using ImageJ software.

### RNA isolation and cDNA synthesis and quantitative PCR

Total RNA was isolated from cells using TRIzol (Thermo Fisher Scientific, #15596018) according to the manufacturer protocol. 1 μg of total RNA was treated with DNase I (#M0303) and reverse transcribed using SuperScript III First-Strand cDNA synthesis kit (Thermo Fisher Scientific, #18080051). The resulting cDNAs were quantified using the real time PCR (StepOne Plus, Thermo Fisher Scientific) to check the expression of autophagy-responsive gene expression with the primer set (Supplementary Table S1). The qPCR fold change was calculated using the ΔΔ*C*_t_ method after normalizing with loading control actin RNA or GAPDH (Supplementary Table S1).

### Anchorage-independent growth assay

Soft agar colony formation assay was performed as described previously(17). Briefly, the bottom layer of soft agar was prepared by mixing equal volume of 1% sterile noble agar, which was maintained at 40°C (Sigma-Aldrich, catalog no. A5431) and 2X growth medium and kept at room temperature for 30 min to solidify. For the top layer, 1×10^4^ cells were resuspended with equal volume of 2X growth medium and 0.6% soft agar, which was also maintained at 40°C and added dropwise on the bottom agar layer. The plates were allowed to solidity for additional 30 min at room temperature. The plates were then kept at humidified chamber at 37°C in the presence of 5% CO_2_ for an additional 3-4 weeks. The visible colonies were captured using 4X bright field objective. 10 random microscopic fields were used for quantification. ImageJ software was used for calculating the colony number and size.

### Measurement of AMP Concentration

AMP concentration was measured according to the manufacturer protocol (BioVision #K229). Briefly, 2X10^7^ cells from WT and DRAIC KO LNCaP cells were harvested either treated with Bay11-7082 or knocking down p65 with siRNA. The cells pellet was washed with 1X PBS, pH 7.4 and lysed with AMP assay buffer and centrifuged at 10,000Xg for 10 min. The protein concentration was measured, and equal amount of protein was used for the assay. The absorbance of the colored product was measured at 570 nm. The AMP concentration of the WT and DRAIC KO cells were measured from the AMP standard curve and expressed as nmol/mg.

### Mouse xenograft

The effect of DRAIC overexpression on tumor growth *in vivo* was validated by intracranial mouse xenograft model. Six-week-old athymic nude mice were procured from The Jackson Laboratory and mice experiments were carried out according to the University of Virginia institutional guidelines. A total of 2×10^5^ U87 cells stably transfected with either EV or DRAIC long noncoding RNA were stereotactically implanted into the striata of immunodeficient mice. After three weeks of injection, mice were subjected to MRI imaging. Tumor volume was calculated according to the published and established protocols.

### DRAIC Exon 2-4 knockout by CRISPR/Cas9

DRAIC KO LNCaP cells were prepared by CRISPR/Cas9 system according to previous publication (17). Briefly, sgRNAs targeting DRAIC exon 2 and 4 were designed using http://crispr.mit.edu/. The sgRNAs were annealed and ligated with quick ligation mixture into px333 vector, which was digested separately with Bbs1 and Bsa1 restriction enzymes respectively. The LNCaP cells were transfected with px333 plasmid containing both the sgRNAs targeting DRAIC exon 2 and exon 4 and selected with the 500 ug/ml antibiotic marker G418. The selected heterogenous resistant population were plated into 96 well plate for single cell expansion. The genomic DNA was isolated from each single clone using the quick genomic DNA isolation kit and DRAIC genomic deletion was confirmed both by PCR and Sanger sequencing.

### Statistical analysis

All data are expressed as mean ± SD from indicated numbers of measurements. The significance was calculated by Student *t* test (paired test, two sided). The differences are called statistically significant if the *P* value is <0.05.

## Supplementary Figure Legends

**Figure S1:**
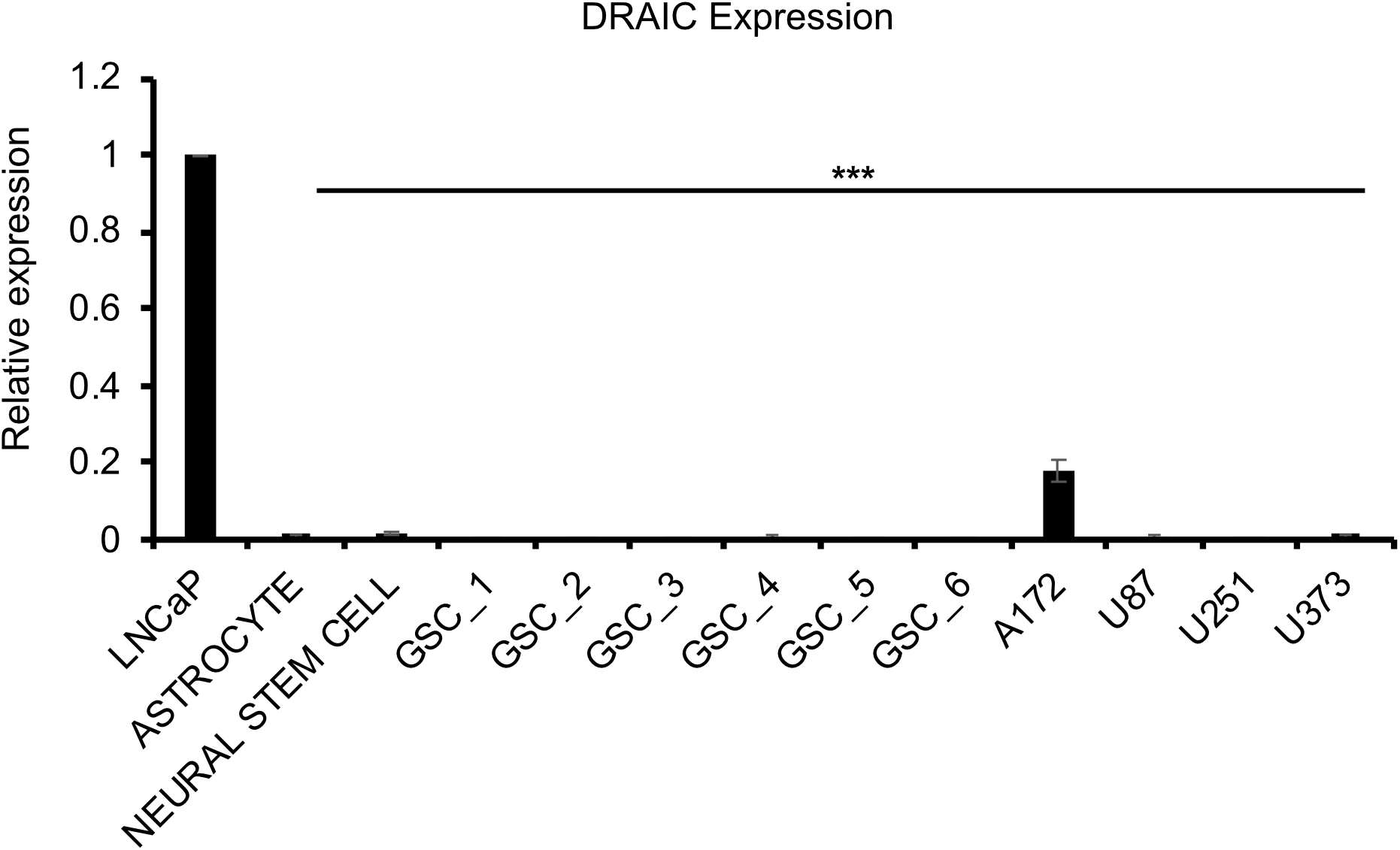
Endogenous DRAIC expression is low in glioma cell lines: The expression of DRAIC is measured in a panel of glioblastoma cell lines and glioma stem cells derived from patients by RT-qPCR. GAPDH is used as an internal control for RT-qPCR normalization. The DRAIC expression in LNCaP cells is used as 1 for comparison. Data are presented as mean ± SD, ***, p<0.001.

**Figure S2:**
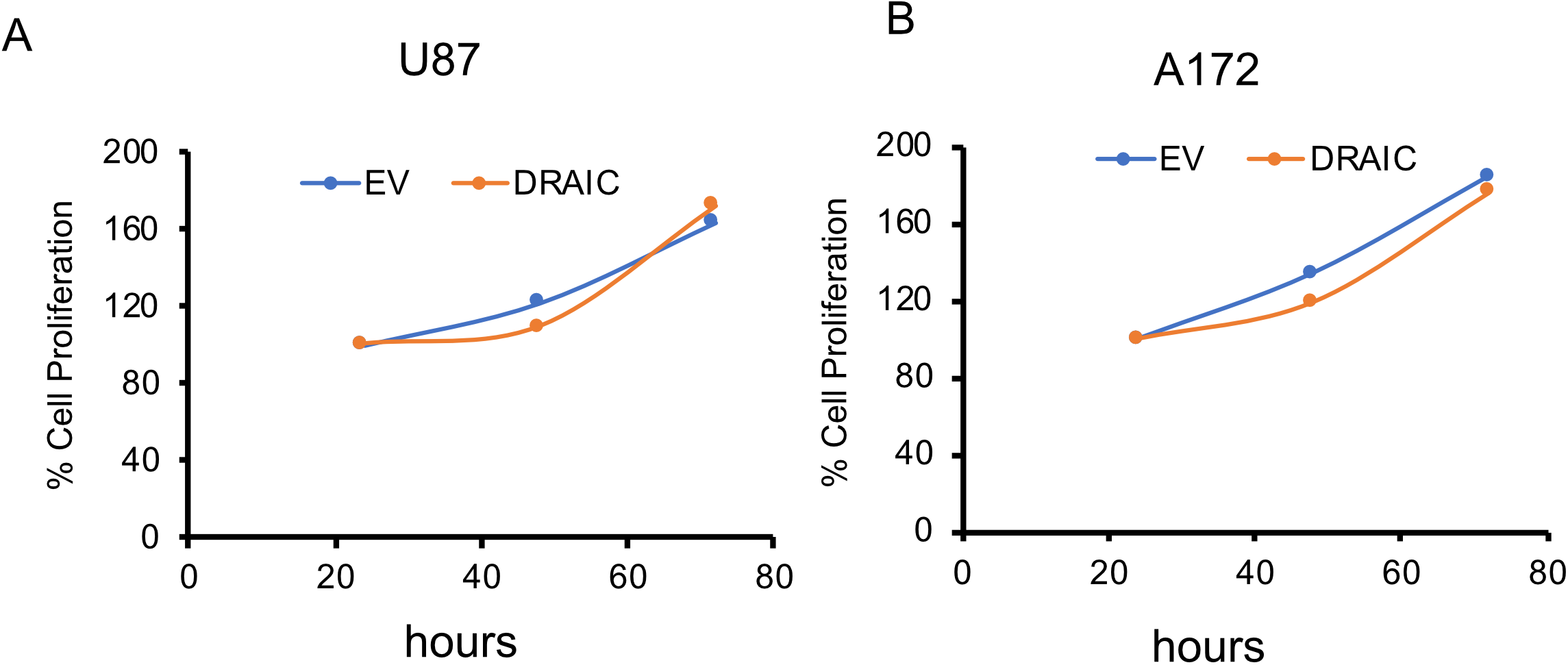
Proliferation of cells adherent on plastic was unchanged when DRAIC is overexpressed in glioblastoma cells: **A-B)** U87 **(A)** and A172 **(B)** cells transfected either with EV or DRAIC and MTT assay was performed at different time points.

**Figure S3:**
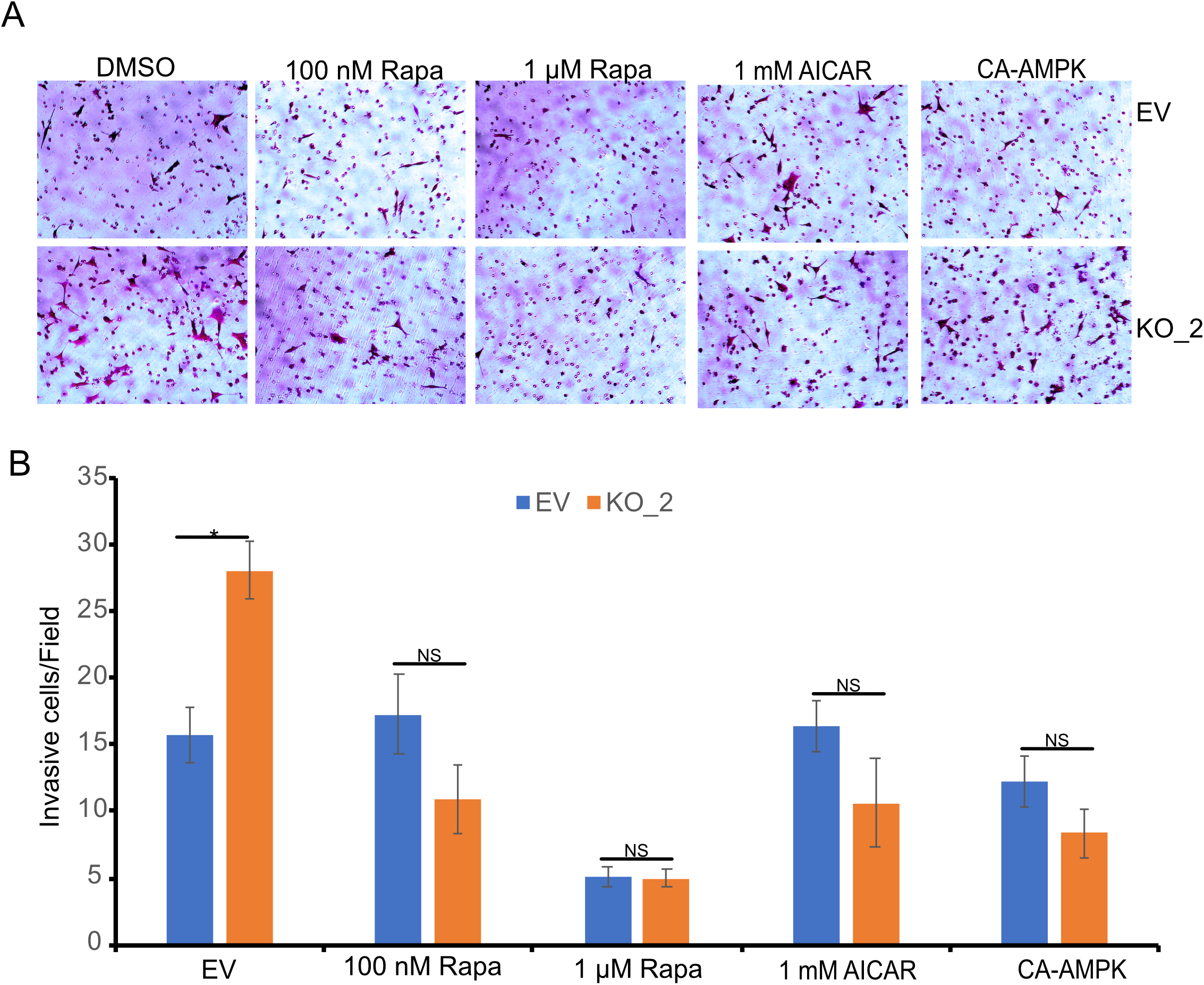
DRAIC KO induced invasion can be reversed by AMPK activators: **A-B)** WT and DRAIC KO LNCaP cells were transfected with either EV or constitutive active AMPK or pre-treated cells with 0.1 and 1 μM rapamycin or 1 mM AMPK activator AICAR for 24 hrs and Matrigel Boyden chamber invasion assay was performed **(A)**. The invaded cells were stained with 0.5% crystal violet and imaged with microscope. 10 random fields were selected for counting **(B)**. Data are presented as mean ± SD *, p<0.05.

**Figure S4:**
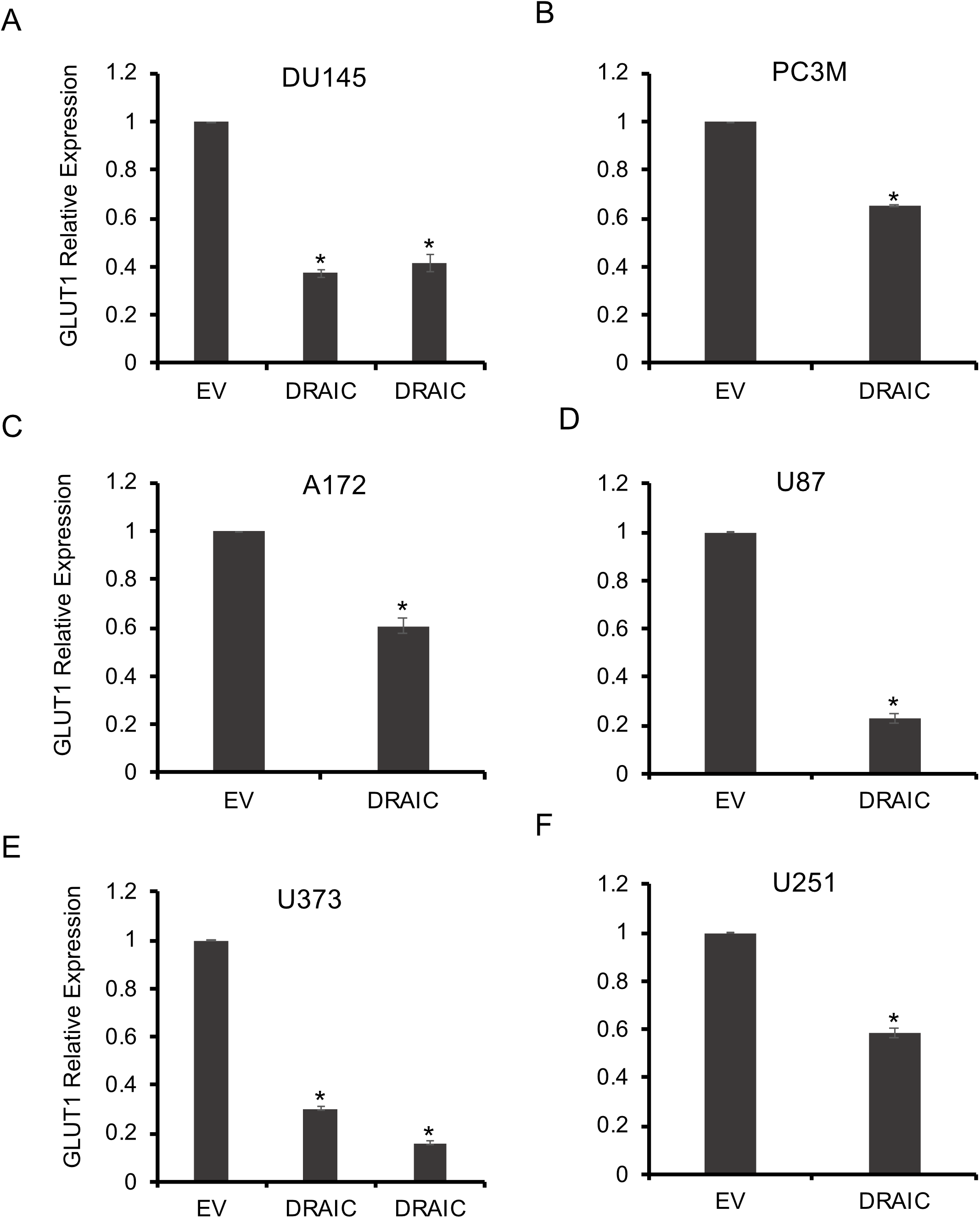
DRAIC overexpression decreases GLUT1 mRNA expression in different cell lines: **A-F)** GLUT1 mRNA level is measured by RT-qPCR in prostate cancer cell lines DU145 **(A)** and PC3M **(B)** cells and glioblastoma cell lines, A172 **(C)**, U87 **(D)**, U373 **(E)**, U251 **(F)**. Data are presented as mean ± SD, *, p<0.05.

**Supplementary Table 1:** Q-RT-PCR primers used in this paper.

## References

1. Demuth T, Berens ME. Molecular mechanisms of glioma cell migration and invasion. Journal of neuro-oncology 2004;70:217–28

2. Urbańska K, Sokołowska J, Szmidt M, Sysa P. Glioblastoma multiforme–an overview. Contemporary oncology 2014;18:307

3. Wen PY, Weller M, Lee EQ, Alexander BM, Barnholtz-Sloan JS, Barthel FP, et al. Glioblastoma in adults: a Society for Neuro-Oncology (SNO) and European Society of Neuro-Oncology (EANO) consensus review on current management and future directions. Neuro-oncology 2020;22:1073–113

4. Anjum K, Shagufta BI, Abbas SQ, Patel S, Khan I, Shah SAA, et al. Current status and future therapeutic perspectives of glioblastoma multiforme (GBM) therapy: A review. Biomedicine & Pharmacotherapy 2017;92:681–9

5. Raizer JJ, Abrey LE, Lassman AB, Chang SM, Lamborn KR, Kuhn JG, et al. A phase II trial of erlotinib in patients with recurrent malignant gliomas and nonprogressive glioblastoma multiforme postradiation therapy. Neuro-oncology 2010;12:95–103

6. Adamson C, Kanu OO, Mehta AI, Di C, Lin N, Mattox AK, et al. Glioblastoma multiforme: a review of where we have been and where we are going. Expert opinion on investigational drugs 2009;18:1061–83

7. Gutschner T, Diederichs S. The hallmarks of cancer: a long non-coding RNA point of view. RNA biology 2012;9:703–19

8. Ransohoff JD, Wei Y, Khavari PA. The functions and unique features of long intergenic non-coding RNA. Nature reviews Molecular cell biology 2018;19:143

9. Bhan A, Mandal SS. Long noncoding RNAs: emerging stars in gene regulation, epigenetics and human disease. ChemMedChem 2014;9:1932–56

10. Hung T, Chang HY. Long noncoding RNA in genome regulation: prospects and mechanisms. RNA biology 2010;7:582–5

11. Statello L, Guo C-J, Chen L-L, Huarte M. Gene regulation by long non-coding RNAs and its biological functions. Nature Reviews Molecular Cell Biology 2020:1–23

12. Yoon J-H, Abdelmohsen K, Gorospe M. Posttranscriptional gene regulation by long noncoding RNA. Journal of molecular biology 2013;425:3723–30

13. Gupta RA, Shah N, Wang KC, Kim J, Horlings HM, Wong DJ, et al. Long non-coding RNA HOTAIR reprograms chromatin state to promote cancer metastasis. Nature 2010;464:1071–6

14. Malek E, Jagannathan S, Driscoll JJ. Correlation of long non-coding RNA expression with metastasis, drug resistance and clinical outcome in cancer. Oncotarget 2014;5:8027

15. Sanchez Calle A, Kawamura Y, Yamamoto Y, Takeshita F, Ochiya T. Emerging roles of long non-coding RNA in cancer. Cancer science 2018;109:2093–100

16. Hajjari M, Salavaty A. HOTAIR: an oncogenic long non-coding RNA in different cancers. Cancer biology & medicine 2015;12:1

17. Saha S, Kiran M, Kuscu C, Chatrath A, Wotton D, Mayo MW, et al. Long noncoding RNA DRAIC inhibits prostate cancer progression by interacting with IKK to inhibit NF-κB activation. Cancer research 2020;80:950–63

18. Sakurai K, Reon BJ, Anaya J, Dutta A. The lncRNA DRAIC/PCAT29 locus constitutes a tumor-suppressive nexus. Molecular cancer research 2015;13:828–38

19. Wang X, Proud CG. The mTOR pathway in the control of protein synthesis. Physiology 2006;21:362–9

20. Holz MK, Ballif BA, Gygi SP, Blenis J. mTOR and S6K1 mediate assembly of the translation preinitiation complex through dynamic protein interchange and ordered phosphorylation events. Cell 2005;123:569–80

21. Easton J, Houghton P. mTOR and cancer therapy. Oncogene 2006;25:6436–46

22. Jung CH, Ro S-H, Cao J, Otto NM, Kim D-H. mTOR regulation of autophagy. FEBS letters 2010;584:1287–95

23. Kim YC, Guan K-L. mTOR: a pharmacologic target for autophagy regulation. The Journal of clinical investigation 2015;125:25–32

24. Egan D, Kim J, Shaw RJ, Guan K-L. The autophagy initiating kinase ULK1 is regulated via opposing phosphorylation by AMPK and mTOR. Autophagy 2011;7:643–4

25. White E. Deconvoluting the context-dependent role for autophagy in cancer. Nature reviews cancer 2012;12:401–10

26. Levy JMM, Towers CG, Thorburn A. Targeting autophagy in cancer. Nature Reviews Cancer 2017;17:528–42

27. Chittaranjan S, Bortnik S, Dragowska WH, Xu J, Abeysundara N, Leung A, et al. Autophagy inhibition augments the anticancer effects of epirubicin treatment in anthracycline-sensitive and-resistant triple-negative breast cancer. Clinical Cancer Research 2014;20:3159–73

28. Taylor MA, Das BC, Ray SK. Targeting autophagy for combating chemoresistance and radioresistance in glioblastoma. Apoptosis 2018;23:563–75

29. Jiang J, Zhang L, Chen H, Lei Y, Zhang T, Wang Y, et al. Regorafenib induces lethal autophagy arrest by stabilizing PSAT1 in glioblastoma. Autophagy 2020;16:106–22

30. Gwinn DM, Shackelford DB, Egan DF, Mihaylova MM, Mery A, Vasquez DS, et al. AMPK phosphorylation of raptor mediates a metabolic checkpoint. Molecular cell 2008;30:214–26

31. Hardie DG, Ashford ML. AMPK: regulating energy balance at the cellular and whole body levels. Physiology 2014;29:99–107

32. Xiao B, Sanders MJ, Underwood E, Heath R, Mayer FV, Carmena D, et al. Structure of mammalian AMPK and its regulation by ADP. Nature 2011;472:230–3

33. Alers S, Löffler AS, Wesselborg S, Stork B. Role of AMPK-mTOR-Ulk1/2 in the regulation of autophagy: cross talk, shortcuts, and feedbacks. Molecular and cellular biology 2012;32:2–11

34. Kim J, Kundu M, Viollet B, Guan K-L. AMPK and mTOR regulate autophagy through direct phosphorylation of Ulk1. Nature cell biology 2011;13:132–41

35. Inoki K, Zhu T, Guan K-L. TSC2 mediates cellular energy response to control cell growth and survival. Cell 2003;115:577–90

36. Egan DF, Shackelford DB, Mihaylova MM, Gelino S, Kohnz RA, Mair W, et al. Phosphorylation of ULK1 (hATG1) by AMP-activated protein kinase connects energy sensing to mitophagy. Science 2011;331:456–61

37. Sanchez AM, Csibi A, Raibon A, Cornille K, Gay S, Bernardi H, et al. AMPK promotes skeletal muscle autophagy through activation of forkhead FoxO3a and interaction with Ulk1. Journal of cellular biochemistry 2012;113:695–710

38. Mair DB, Ames HM, Li R. Mechanisms of invasion and motility of high-grade gliomas in the brain. Molecular biology of the cell 2018;29:2509–15

39. Guertin DA, Sabatini DM. Defining the role of mTOR in cancer. Cancer cell 2007;12:9–22

40. Pópulo H, Lopes JM, Soares P. The mTOR signalling pathway in human cancer. International journal of molecular sciences 2012;13:1886–918

41. Sarbassov DD, Ali SM, Sabatini DM. Growing roles for the mTOR pathway. Current opinion in cell biology 2005;17:596–603

42. Shaw RJ. LKB1 and AMP-activated protein kinase control of mTOR signalling and growth. Acta physiologica 2009;196:65–80

43. Xu J, Ji J, Yan X-H. Cross-talk between AMPK and mTOR in regulating energy balance. Critical reviews in food science and nutrition 2012;52:373–81

44. Xia Y, Shen S, Verma IM. NF-κB, an active player in human cancers. Cancer immunology research 2014;2:823–30

45. Wang X, Liu R, Qu X, Yu H, Chu H, Zhang Y, et al. α-Ketoglutarate-activated NF-κB signaling promotes compensatory glucose uptake and brain tumor development. Molecular cell 2019;76:148–62. e7

46. Dan HC, Cooper MJ, Cogswell PC, Duncan JA, Ting JP-Y, Baldwin AS. Akt-dependent regulation of NF-κB is controlled by mTOR and Raptor in association with IKK. Genes & development 2008;22:1490–500

47. Wu ZR, Yan L, Liu YT, Cao L, Guo YH, Zhang Y, et al. Inhibition of mTORC1 by lncRNA H19 via disrupting 4E-BP1/Raptor interaction in pituitary tumours. Nature communications 2018;9:1–14

48. Malakar P, Shilo A, Mogilevsky A, Stein I, Pikarsky E, Nevo Y, et al. Long noncoding RNA MALAT1 promotes hepatocellular carcinoma development by SRSF1 upregulation and mTOR activation. Cancer research 2017;77:1155–67

49. Chen J-F, Wu P, Xia R, Yang J, Huo X-Y, Gu D-Y, et al. STAT3-induced lncRNA HAGLROS overexpression contributes to the malignant progression of gastric cancer cells via mTOR signal-mediated inhibition of autophagy. Molecular cancer 2018;17:1–16

50. Du Y, Wang L, Chen S, Liu Y, Zhao Y. lncRNA DLEU1 contributes to tumorigenesis and development of endometrial carcinoma by targeting mTOR. Molecular carcinogenesis 2018;57:1191–200

51. Li Z, Yang Z, Passaniti A, Lapidus RG, Liu X, Cullen KJ, et al. A positive feedback loop involving EGFR/Akt/mTORC1 and IKK/NF-κB regulates head and neck squamous cell carcinoma proliferation. Oncotarget 2016;7:31892

52. Xue J, Zhong S, Sun B-m, Sun Q-F, Hu L-Y, Pan S-J. Lnc-THOR silencing inhibits human glioma cell survival by activating MAGEA6-AMPK signaling. Cell death & disease 2019;10:1–13

53. Sun M, Gadad SS, Kim D-S, Kraus WL. Discovery, annotation, and functional analysis of long noncoding RNAs controlling cell-cycle gene expression and proliferation in breast cancer cells. Molecular cell 2015;59:698–711

54. Tiessen I, Abildgaard MH, Lubas M, Gylling HM, Steinhauer C, Pietras EJ, et al. A high-throughput screen identifies the long non-coding RNA DRAIC as a regulator of autophagy. Oncogene 2019;38:5127–41

55. Zhao D, Dong J-T. Upregulation of long non-coding RNA DRAIC correlates with adverse features of breast cancer. Non-coding RNA 2018;4:39

